# Memory in Repetitive Protein–Protein Interaction Series – in Memory of the Late Professor Robert M. Nerem

**DOI:** 10.1101/2022.10.01.510459

**Authors:** Aaron M. Rosado, Yan Zhang, Hyun-Kyu Choi, Samuel M. Ehrlich, Fengzhi Jin, Arash Grakoui, Brian D. Evavold, Cheng Zhu

**Author notes:** These authors contributed equally (Aaron M. Rosado, Yan Zhang). 95 Zhengtong Road, Building No. 3, Apt #1102, Shanghai, China (Yan Zhang). Department of Pathology, University of Utah School of Medicine, Salt Lake City, UT, USA.

## Abstract

Over the past three decades, the senior author had interacted with and been mentored by the late Professor Robert M. Nerem. In his memory, the authors summarized several observations made, ideas conceptualized, and mathematical models developed during this period for quantitatively analyzing memory effects in repetitive protein–protein interactions (PPI). Interactions between proteins in an organism coordinate its biological processes and may impact its responses to changing environment and diseases through feedback systems. Feedback systems function by using changes in the past to influence behaviors in the future, which we refer here as memory. Specifically, we consider how proteins on cell or in isolation retain information about prior interactions to impact current interactions. The micropipette, biomembrane force probe and atomic force microscopic techniques were used to repeatedly assay several PPIs. The resulting time series were analyzed by a previous and two new models to extract three memory indices of short (seconds), intermediate (minutes), and long (hours) timescales. We found that interactions of cell membrane, but not soluble, T cell receptor (TCR) with peptide-major histocompatibility complex (pMHC) exhibits short-term memory that impacts on-rate, but not off-rate of the binding kinetics. Peptide dissociation from MHC resulted in intermediate- and long-term memories in TCR–pMHC interactions. However, we observed no changes in kinetic parameters by repetitive measurements on living cells over intermediate timescale using stable pMHCs. Parameters quantifying memory effects in PPIs could provide additional information regarding biological mechanisms. The methods developed herein also provide tools for future research.

## Introduction

The corresponding author joined the Bioengineering faculty of Georgia Institute of Technology led by the late Professor Robert M. Nerem in 1990 as a mathematical modeler. Under the support and mentorship of the late Professor Nerem, he developed a series of experimental capabilities in his laboratory with his students over the past three decades, including the micropipette (MP)^1^ (Fig. 1A), biomembrane force probe (BFP)^2, 3^ (Fig. 1B), and atomic force microscopy (AFM)^4, 5^ (Fig. 1C). A major method developed by the Zhu lab is the adhesion frequency assay that combines the ability of MT, BFP and AFM to manipulate interactions and measure adhesive forces between two surfaces with a mathematical model for probabilistic kinetics of a small number of receptor–ligand interaction.^1, 6, 7^ This assay represents a single-cell and even single-molecule approach designed for measuring *in situ* receptor–ligand interaction across the interface between two cells. Not only does it possess a much higher sensitivity than many of the existing approaches for characterizing receptor–ligand interaction kinetics and affinity, but it can also integrate information about molecular and cellular level changes in biological systems.^8, 9^ Collaborating with colleagues at Emory University and other institutions, including two co-authors of this paper, the Zhu lab employed the adhesion frequency assay to study ligand binding by a wide variety of receptors, including Fc γ receptors (FcγRs),^10^ T cell receptors (TCRs),^11-15^ and platelet glycoprotein Iα (GPIbα).^16-18^

**Figure 1.**
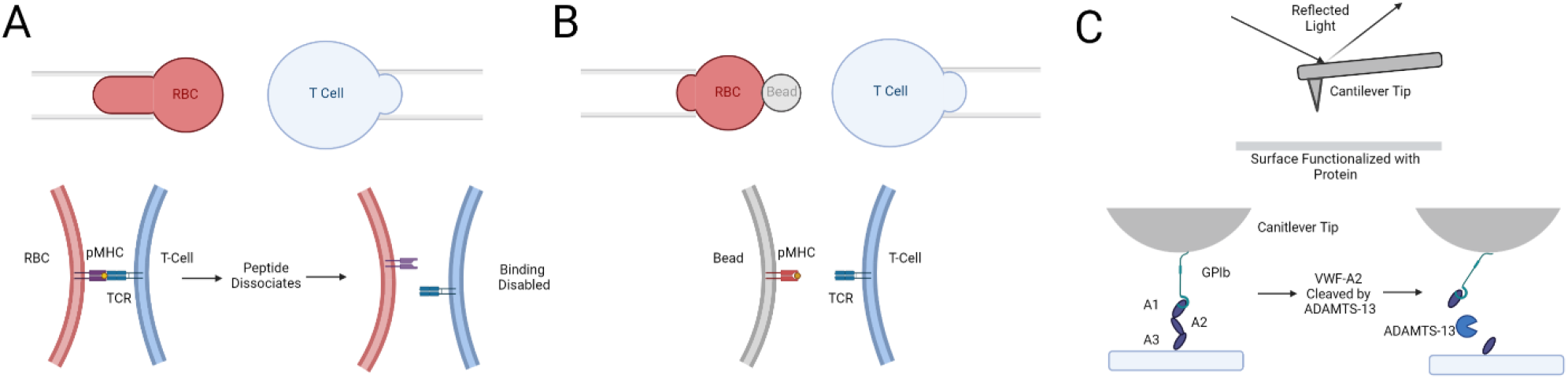
Schematics of the three ultrasensitive force transducers employed (*top*) and the molecular systems used (*bottom*) to perform the adhesion frequency assay. *A*. Micropipette aspirated RBC bearing p:H2-K^b^α3A2 or p:I-E^k^ (*left*) testing against an OT1 or 3.L2 T cell (*right*). *B*. Biomembrane force probe (BFP) bearing gp33:H2-D^b^α3A2 or TPI:HLA-DR1 (*left*) testing against a P14 T cell or Jurkat cell expressing or E8 TCR (*right*). In some experiments, the BFP was coated with E8 TCR to test against a THP-1 cell expressing TPI:HLA-DR1. *C*. Atomic force microscopy with a GPIbα-bearing cantilever tip (*upper*) testing against VWR-A1A2A3 tri-domain coated surface (*lower*). Also depicted are two mechanisms of causing memory effects of intermediate and long timescales, peptide dissociation (A) and proteolytic cleavage (C).

Data from the adhesion frequency assay are unique in their ability to bridge knowledge about structure and function of proteins and cells.^10, 13-15, 19-27^ Despite these advantages and utilities, such data remain underanalyzed and underutilized. In particular, the original analysis of the adhesion frequency assay treats the experimentally generated binary adhesion score sequence as a Bernoulli process of an independent and identically distributed (i.i.d.) random variable neglecting any relationship between past and current interactions, i.e., assumes no memory. In many biological systems, however, prior molecular interactions can and do influence subsequent interactions. The relationship between prior and current interactions may reveal feedback mechanisms that could be important to many properties of the biological systems in question, such as sensitivity, stability, robustness, adaptiveness, resilience, etc., even at molecular and cellular levels. As an initial step towards developing methods for characterizing feedback systems, we built a model for analyzing the impact on current interaction by immediate past’s outcome, i.e., short-term memory.^28^ Within the context of this work, we refer to the idea that molecular and cellular systems retain information about past molecular interactions as memory. Memory can manifest as both irreversible and reversible changes that influence subsequent interactions and may reflect mechanisms that occur in different time scales ranging from seconds to hours. At a molecular level, memory can involve interactions between proteins mediating reversible changes like phosphorylation and de-phosphorylation and irreversible changes such as proteolytic cleavage. At a cellular level, memory can be caused by changes in receptor expression through internalization, degradation and recycling, as well as proteolytic shedding.

In dedication to the memory of the late Professor Robert M. Nerem, here we extend our previous memory analysis^28^ to include models of memory effects in three different timescales, ranging from seconds, minutes, to hours. We present unpublished models for the analyses of memory effects in the intermediate and long timescales. We apply these three memory models to a wide range of data collected over decades by different students and postdoctoral scholars of the Zhu lab, focusing on interactions of TCR with peptide-major histocompatibility complex (pMHC) but also including those of Fc γ receptor IIIa (FcγRIIIa) with IgG Fc and anti-FcγRIIIa, and GPIbα with von Willebrand factor (VWF). We extend the short-term memory analysis from adhesion probability to bond lifetime under a range of forces, which are related to binding affinity and force-dependent off-rate of dissociation. Our results demonstrate the validity and utility of the models, which provide analytical tools to classify and organize data as well as extract quantitative information in the forms of three memory indices. Future studies will apply these models to more systems and relate the memory indices to biological functionalities and mechanisms.

### Model development

We used the adhesion frequency assay^1, 6, 7^ to measure *in situ* kinetics of cross-junctional interactions between two surfaces respectively expressing receptors and ligands. One surface is part of a force transducer that enables detection of tensile forces between the two surfaces as a mechanical method to detect adhesion. Three types of force transducers employed in this study include: 1) a human red blood cell (RBC) pressurized by MT aspiration,^1, 11, 29^ 2) a glass bead attached to the pressurized RBC, or BFP,^13, 30^ and 3) an AFM^4, 18, 31^ (Fig. 1A-C, see Methods). These ultrasensitive transducers have a single piconewton (pN) force sensitivity, which detect adhesion mediated by as low as a single receptor–ligand bond.^6^ The adhesion frequency assay used a series of repeated contact-retraction cycles between a single-paired receptor-bearing surface and force transducer to generate a binary sequence of positive or negative outcomes (1 for adhesion and 0 for no adhesion), which contain memory information in both short and intermediate timescales.

#### Model for no memory

The simplest binary adhesion score sequence analysis model the repeated contact-retraction cycles as a Bernoulli process satisfying the i.i.d. assumption that implies no memory. Let the probability of having the positive outcome (binding, score = 1) be *p* so the probability of having the negative outcome (no binding, score = 0) be 1 – *p. p* can be estimated from the average of all adhesion scores in the sequence, which is the adhesion frequency *P*_a_. In our experiments, the ligand site densities were adjusted to such a range that the adhesion frequency would be 15% < *P*_a_ < 85%. This condition ensures binding to be mediated by a low number of receptor–ligand bonds for the Poisson distribution of bonds to apply.^1^ The Poisson distribution depends on only one parameter, <*n*>, the average number of bonds per contact that relates to the probability of having no bond by 1 – *p* = exp(– <*n*>). By modeling the receptor–ligand interaction as a second-order forward (driven by the densities of receptors, *m*_r_, and ligands, *m*_l_, via mass action) and first-order reverse (driven by the average number of bonds, <*n*>) reversible reaction, *P*_a_ can be related to experimentally controlled and measured variables by:^1, 6^

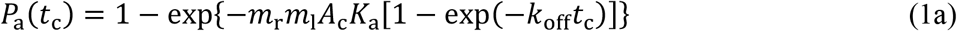

where *A*_c_ (kept constant throughout) and *t*_c_ (kept constant for a sequence of repeated contact-retraction cycles) are the contact area and contact duration. *K*_a_ is two-dimensional (2D) affinity (with 2D area units, e.g., µm^2^, rather than a 3D volume) and *k*_off_ is the off-rate (in s^-1^). The 2D on-rate can be calculated as *k*_on_ = *K*_a_ × *k*_off_ (in µm^2^s^-1^). The term behind the minus sign inside the curly brackets on the right-hand side of Eq. 1 can be identified as the average number of bonds <*n*>, i.e.,

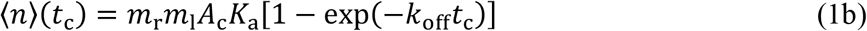

given that *P*_a_ = *p* = 1 –exp(– <*n*>) under the current assumption of no memory.

#### Model for short memory

A close inspection of many binary adhesion score sequences suggests that the i.i.d. assumption may not be always valid as these sequences sometimes exhibit non-Bernoulli behaviors. The models used to characterize these non-Bernoulli behaviors depend on the timescale involved. For example, the i.i.d. assumption would be violated if the probability *p* of having a current contact-retraction test positive outcome (binding) becomes dependent on previous test outcomes meaning the present has memory of the past. The one-step Markov process models the simplest deviation from the Bernoulli process, where the positive outcome of each test has different probabilities, *p* + Δ*p* or *p*, depending on whether the outcome of the immediate past test is positive or negative. Here Δ*p* is termed short-term memory index and can have a value >, < or = 0, corresponding to three scenarios of adhesion in the past that 1) enhances (positive memory), 2) inhibits (negative memory), or 3) has no influence on (no memory) future adhesions. The memory effect quantified by Δ*p* has a very short timescale because each contact-retraction cycle takes no more than a couple of seconds cycling time plus the contact duration. Another reason for the short time scale is that the one-step Markov process considers memory only of the immediate past.

We previously developed a set of models for such short memory effects.^28, 32, 33^ The memory index Δ*p* can be evaluated using two methods, direct calculation or fitting to adhesion cluster distribution.^28^ The direct calculation is based on the definitions:

1. *p* + Δ*p* is the conditional probability of positive outcome of the present contact-retraction test given that the outcome of the immediate past test is positive;
2. *p* is the conditional probability of positive outcome of the present test given that the outcome of the immediate past test is negative;
3. 1 – (*p* + Δ*p*) is the conditional probability of negative outcome of the present test given that the outcome of the immediate past test is positive; and
4. 1 – *p* is the conditional probability of negative outcome of the present test, given that the outcome of the immediate past test is negative.

From the adhesion score sequences measured from a given biological system, we segregated the binding events into four types (cf. Fig. 3A) and enumerated each: 1) binding events in the immediate past test of which also are binding events, *n*_11_; 2) binding events the immediate past test of which is a no binding event, *n*_01_; 3) no binding events the immediate past test of which is a binding event, *n*_10_; and 4) no binding events the immediate past test of which also is a no binding event, *n*_00_. Their occurrence frequencies, *n*_11_/(*n*_11_ + *n*_10_), *n*_01_/(*n*_11_ + *n*_10_), *n*_10_/(*n*_01_ + *n*_00_), and *n*_00_/(*n*_01_ + *n*_00_), where *n*_11_ + *n*_10_ = *n*_01_ + *n*_00_ = *n* is the total test events in a particular adhesion test sequence, can be used as estimates of *p* + Δ*p, p*, 1 – (*p* + Δ*p*), and 1 – *p* ^28^. Therefore,

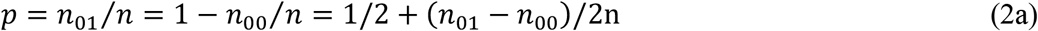

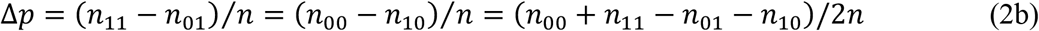

**Figure 2.**
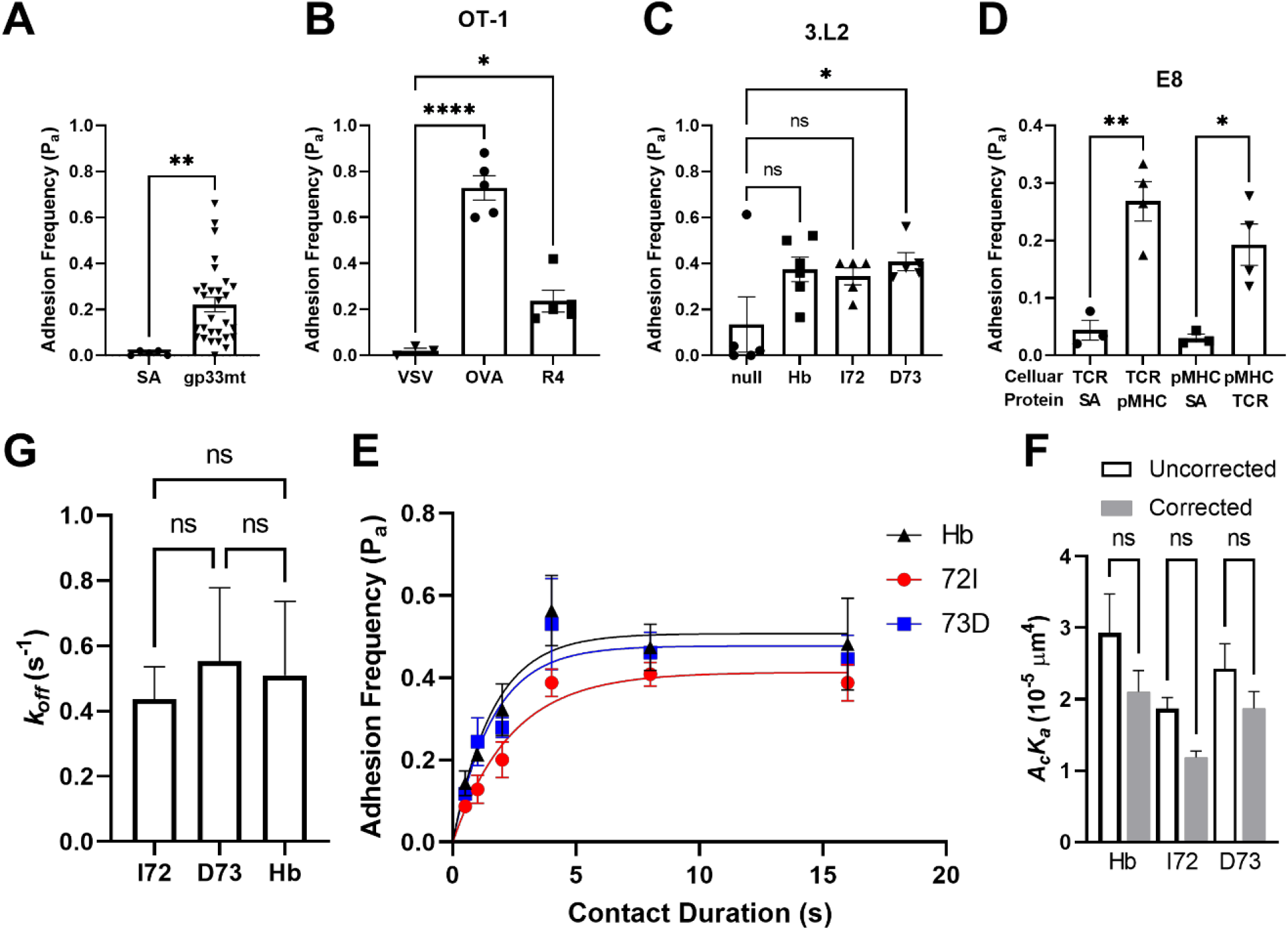
Binding specificity controls and *in situ* kinetic analysis under the assumption of no memory. *A-D*. Steady-state adhesion frequencies between T cells expressing P14 (A), OT1 (B), 3.L2 (C), or E8 TCR interacting with indicated pMHCs coated on force transducer surfaces (BFP probes in A and D or RBCs in B and C) were abolished when the cognate pMHCs were either not coated on the streptavidin (SA) surface (A and D) or replaced by null pMHCs (B and C). Another set of data in D were generated using BFP coated with C-terminally biotinylated E8 TCRαβ ectodomain to test against THP-1 cells expressing TPI-HLA-DR1. Each point was estimated from the average of binary adhesion scores from ≥50 repeated contact-retraction test cycles between a single pair of cell and force transducer (RBC or BFP) and bars ± error bars present mean ± SEM of all points. \**P* = 0.05, \*\**P* = 0.01, \*\*\*\**P* = 0.0001 by one-way ANOVA. *E*. Mean ± SEM adhesion frequency *P*_a_ (n≥5 cell-bead pairs tested 50 times each per point) of CD4^+^ T cells from 3.L2 TCR transgenic mice interacting with indicated peptides presented by mouse class II MHC (I-E^k^) were plotted vs. contact duration *t*_c_. The original model that assumes no memory, Eq. 1a, was fit (curves) to the data (*points*), together with the densities (in *µ*m^-2^) of the TCR (*m*_r_, 200 *µ*m^-2^) and indicated p:I-E^k^ (*m*_l_, 144 *µ*m^-2^) measured by flow cytometry, to evaluate the model parameters. *F, G*. The best-fit values of the effective 2D affinity *A*_c_*K*_a_ (F) and off-rate constant *k*_off_ (G) of the 3.L2 TCR interacting with the indicated pMHC-II, with SEM estimated from the scattering of the data in E by computing the Hessian matrix of the *χ*^2^ function. The gray bars in (F) represents values after the correction of the short-term memory index Δ*p* using data from Fig. 3D.

**Figure 3.**
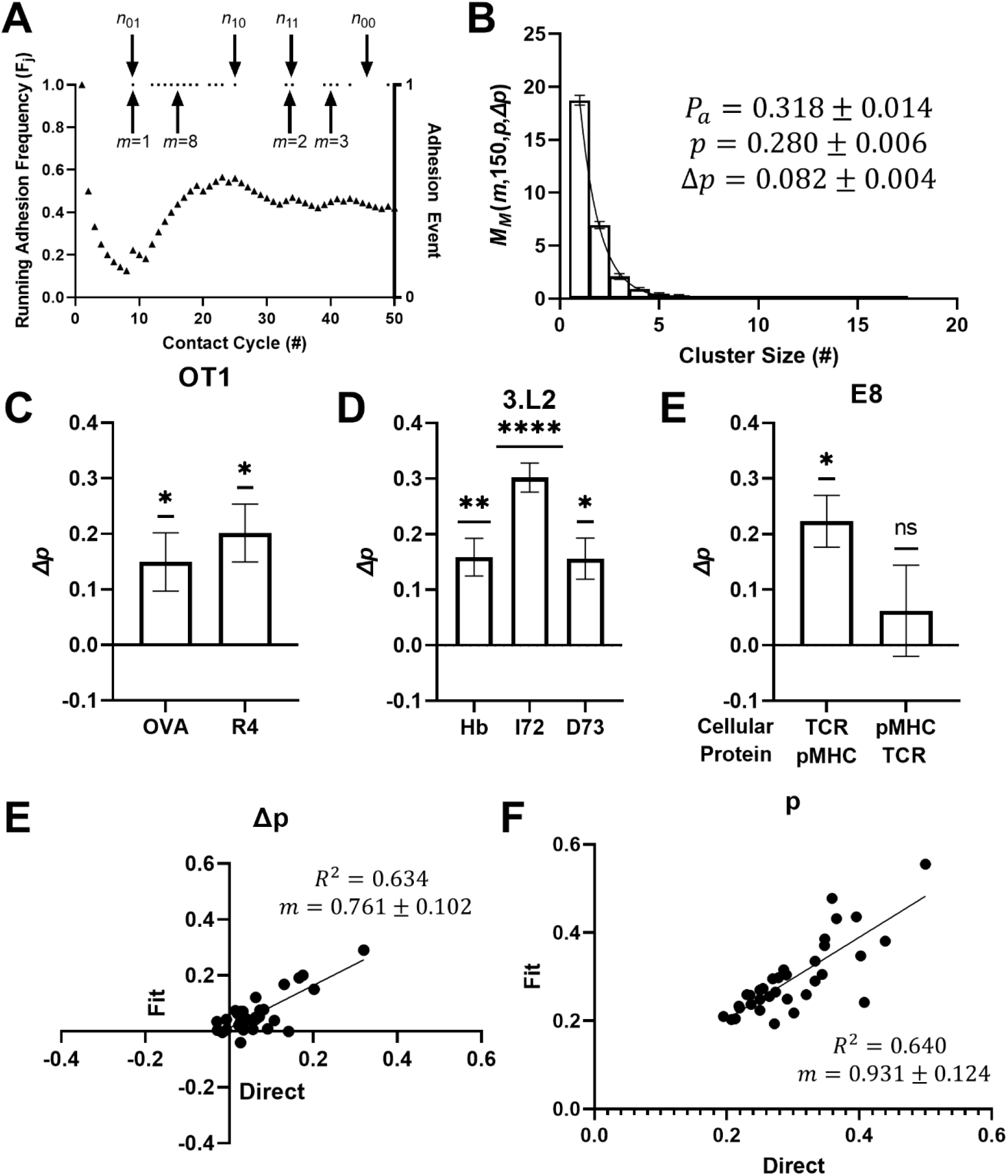
Analysis of memory effect of a short timescale. *A*. Representative running adhesion frequency (left ordinate) and adhesion score (right ordinate, only the positive scores are shown) vs. contact-retraction cycles for the P14 TCR–gp33:H2-D^b^α3A2 interaction. The four types of events *n*_ij_ needed for direct calculation of *p* and Δ*p* using Eq. 2 are indicated. Also indicated are clusters of adhesion events of sizes *m* = 1, 2, 3, and 8. *B*. Representative adhesion cluster size distribution (bars, mean ± SEM) and the model fit by Eq. 3 (curve) for the P14 TCR–gp33:H2-D^b^α3A2 interaction, with best-fit parameters indicated. *C-E*. Mean ± SEM short-term memory index Δ*p*, estimated by the fitting of the cluster size distribution by Eq. 3, for OT1 TCR interacting with indicated pMHC-I ligands (C), 3.L2 TCR interacting with indicated pMHC-II ligands (D), and E8 TCR interacting with TPI:HLA-DR1 in both the normal (cellular TCR vs. soluble pMHC) and inverted (cellular pMHC vs. soluble TCR) configurations (E). ns = no significance, \**P* = 0.05, \*\**P* = 0.01, \*\*\*\**P* = 0.0001 by one-sample t test for comparing if Δ*p* > 0 or not. *F, G*. Comparison between two methods, model fit by Eq. 3 or direct calculation by Eq. 2, for estimating the Δ*p* (E) and *p* (F) values. Each point represents two values estimated using data from a single pair of P14T cell and gp33:H2-D^b^ coated BFP probe by two methods. The line represents linear fit. The slope *m* and *R*^2^ values of the fits are indicated.

The second method to evaluate *p* and Δ*p* from experiment is fitting the experimentally measured to the theoretically expected cluster size distribution:^28^

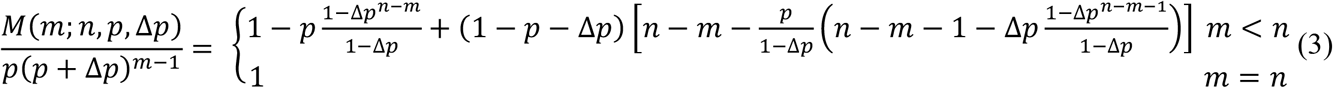

where *m* is the cluster size, i.e., the number of positive outcomes appeared consecutively (cf. Fig. 3A). For a given adhesion test sequence of *n* total events, we can enumerate the frequency *M* of cluster size of *m* (*m* = 1, 2, …, *n*) and fit Eq. 3 to the data to evaluate *p* and Δ*p* as fitting parameters.

Due to the presence of the memory effect, the adhesion frequency *P*_a_ is a function of *p* and Δ*p*:^28^

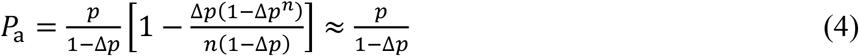

The far-right side of Eq. 4 approximates for large *n* (usually ≥50). When Δ*p* = 0, *P*_a_ = *p*. Note that the average number of bonds <*n*> relates to *p* instead of *P*_a_ regardless of the Δ*p* value, i.e.,

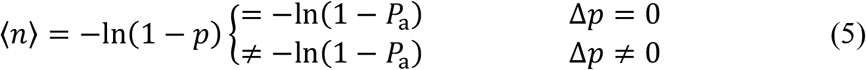

The upper branch of Eq. 5 is identical to Eq. 1 whereas the lower branch corrects for the case when Δ*p* ≠ 0. Note <*n*> = - ln(1 – *p*) is related to the receptor-ligand interaction effective 2D affinity (*A*_c_*K*_a_) and off-rate (*k*_off_) by Eq. 1b.

Therefore, for a given set of adhesion test sequences, we need to first determine from either direct calculation (Eq. 2a) or fitting to adhesion cluster distribution (Eq. 3) to obtain *p* as a function of the receptor density *m*_r_, ligand density *m*_l_, and contact duration *t*_c_ before evaluating for *A*_c_*K*_a_ and *k*_off_ from Eq. 1b)

#### Model for intermediate memory

Let us next consider memory effects of an intermediate timescale of minutes, as exemplified by the cleavage of the von Willebrand factor tri-domain A1A2A3 by the protease ADAMTS13.^31^ The cryptic cleavage site buried inside the folded A2 domain prevents it from being accessed by the proteolytic enzyme ADAMTS13. In the adhesion frequency experiment (Fig. 1C), the A3 domain was immobilized on a cover glass and A1 domain binding by the platelet receptor GPIbα coated on the AFM cantilever tip was tested repeatedly using sequential contact-retraction cycles.^31^ When a binding event was detected by a tensile force signal upon moving the cantilever tip away from the cover glass, the pulling force exerted on the A1A2A3 domain might unfold the A2 domain, exposing the cleavage site. In the presence of ADAMTS13 in solution, the proteolytic enzyme might cut open the peptide bond, breaking the A1A2A3 tri-domain into two polypeptide fragments. This gave rise to a rupture event force signal indistinguishable from a receptor–ligand dissociation event because both events caused a molecular complex rupture physically linking the AFM cantilever tip to the cover-glass. The resulting force signal captured the springing back of the bent cantilever. However, receptor–ligand dissociation is reversible, so the dissociated A1A2A3 tri-domain remained intact allowing for rebinding in the next adhesion test. By comparison, the rupture event was irreversible because the two polypeptide fragments could not rejoin to reform the A1A2A3. As a result, the ligand number available for binding decreased in the next adhesion test, resulting in reduced binding function ^31^. Note that both the force-induced unfolding of the A2 domain and its proteolytic cleavage by ADAMTS13 are stochastic, which may or may not occur in any given test cycle. Hence, the decrease in binding function manifests as diminished adhesion probability over a time sequence containing many contact cycles, giving rise to the memory effect.

In the irreversible process, the present positive test outcome would be affected not only by the outcomes of the immediate past test, but all previous test sequence outcomes. Moreover, if a A1A2A3 tri-domain cleaved at the *j*th test would not be available for binding in not only the (*j* + 1)th test, but also any other tests after that. In other words, binding’s impact is cumulative. Since a full repeated test cycle sequence usually takes several minutes to complete experimentally, such memory effect has an intermediate timescale of minutes. Here we used a phenomenological model for this process type that exhibits a cumulative effect over a progressive running adhesion frequency change:^34^

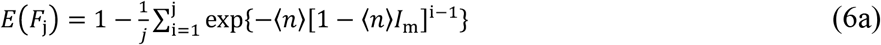

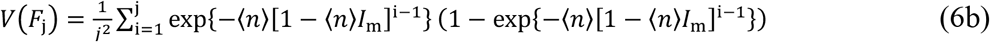

where *F*_j_ is the random variable for the running adhesion frequency over *j* repetitive adhesion tests. *E*(*F*_j_) and *V*(*F*_j_) are the expected average and variance of *F*_j,_ respectively. *I*_m_ is an irreversibility index for measuring the intermediate timescale memory effect. In the absence of irreversibility, *I*_m_ = 0, Eq. 6a and b reduce to:

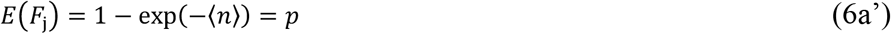

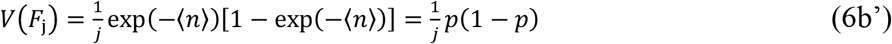

and we recover Bernoulli process properties. Thus, the non-zero *I*_m_ captures the non-Bernoulli effect. The positive and negative *I*_m_ values correspond to two scenarios: adhesions in the past progressively 1) reducing and 2) enhancing future adhesions, respectively.

#### Model for long memory

Finally, let us consider memory effects of a long timescale of hours exemplified by peptide dissociation from micropipette adhesion experiment RBCs coated with MHC molecules (Fig. 1A). After coating and washing, the pMHC molecules on the RBCs exposed to an infinitely dilute solution enables peptide dissociation from the pMHC complex. The first-order irreversible dissociation kinetics models this:

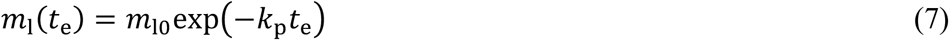

where *k*_p_ is the peptide dissociation rate constant and *t*_e_ is the elapsed time during which peptide dissociation occurs. It follows from Eqs. 1 and 7 that:

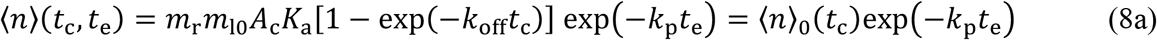

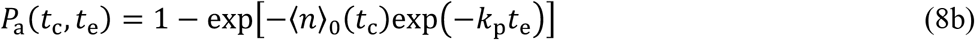

Equation 8 describes an exponential decrease bond formation ability due to ligand function loss over time. Similar scenarios include receptor down-regulation over time, or even upregulation over time due to activation associated with a negative *k*_p_ value. In practice, the experimenter usually uses a sufficiently large *t*_c_ to simplify Eq. 8a and b because exp(–*k*_off_*t*_c_) ⟶ 0 as *t*_c_ ⟶ ∞. The simplified equations are:

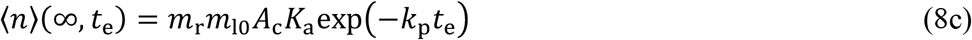

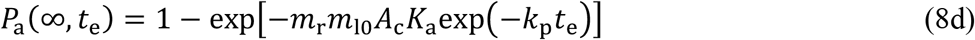

A related parameter is the half time of peptide dissociation, *t*_½_, defined as time required for half peptide dissociation from MHC, i.e., *m*_l_(*t*_½_) = ½ *m*_l0_. It follows from Eq. 7 that *t*_½_ = ln2/*k*_p_.

## Results

### Analysis of in situ kinetics assuming no memory

We used the adhesion frequency assay^1, 6, 7^ to measure *in situ* kinetics of cross-junctional interactions between TCR on T cell surfaces and pMHC-functionalized surrogate antigen presenting cells (APCs). Our experiments employed both CD4^+^ and CD8^+^ T cells from 3.L2, P14, or OT1 TCR transgenic mice, and human E8 TCR expressed on a Jurkat cell line. The surrogate APCs were RBCs for MP experiments (used to test OT1 and 3.L2 T cells) and glass beads for BFP experiments (used to test P14 and E8 T cells) coated with corresponding TCR ligands: p:I-E^k^, p:H2-D^b^α3A2, p:H2-K^b^α3A2, or p:HLA-DR1. We first performed control experiments to test whether measured adhesions were mediated by specific TCR–pMHC interactions. The specificities of the adhesion frequencies to the p:H2-D^b^α3A2 (for P14 TCR, Fig. 2A), p:H2-K^b^α3A2 (for OT1 TCR, Fig. 2B), p:I-E^k^ (for 3L.2 TCR, Fig. 2C), and p:HLA-DR1 (for E8 TCR, Fig. 2D) were confirmed as they were abolished when the TCR or pMHC were either not coated on the BFP probe (SA for P14 and E8) or replaced by null pMHC (VSV for OT1 or MCC for 3.L2).

We previously reported the 2D kinetic parameters of the OT1^13^, 3.L2^11^, P14^15^, and E8^35^ TCRs interacting with their corresponding ligands. Here we analyzed the interactions of 3.L2 TCR with ligands coated on RBCs by the CrCl method different from the biotin-streptavidin method used in previous studies (see Methods). Using a no memory assumption, we plotted the adhesion frequency *P*_a_ vs. contact duration *t*_c_ data and fitted the measured binding curves with Eq. 1a (Figs. 2E and 4G-I). Evidently, the model fits all data very well. Using receptor and ligand densities measured from independent flow cytometry experiment (see Methods) we found the best-fit parameters the effective 2D affinity *A*_c_*K*_a_ and off-rate *k*_off_ for 3.L2 TCR interaction with three peptides, Hb, I72, and D73 presented by mouse pMHC class II, I-E_k_ (Fig. 2F and G).

**Figure 4.**
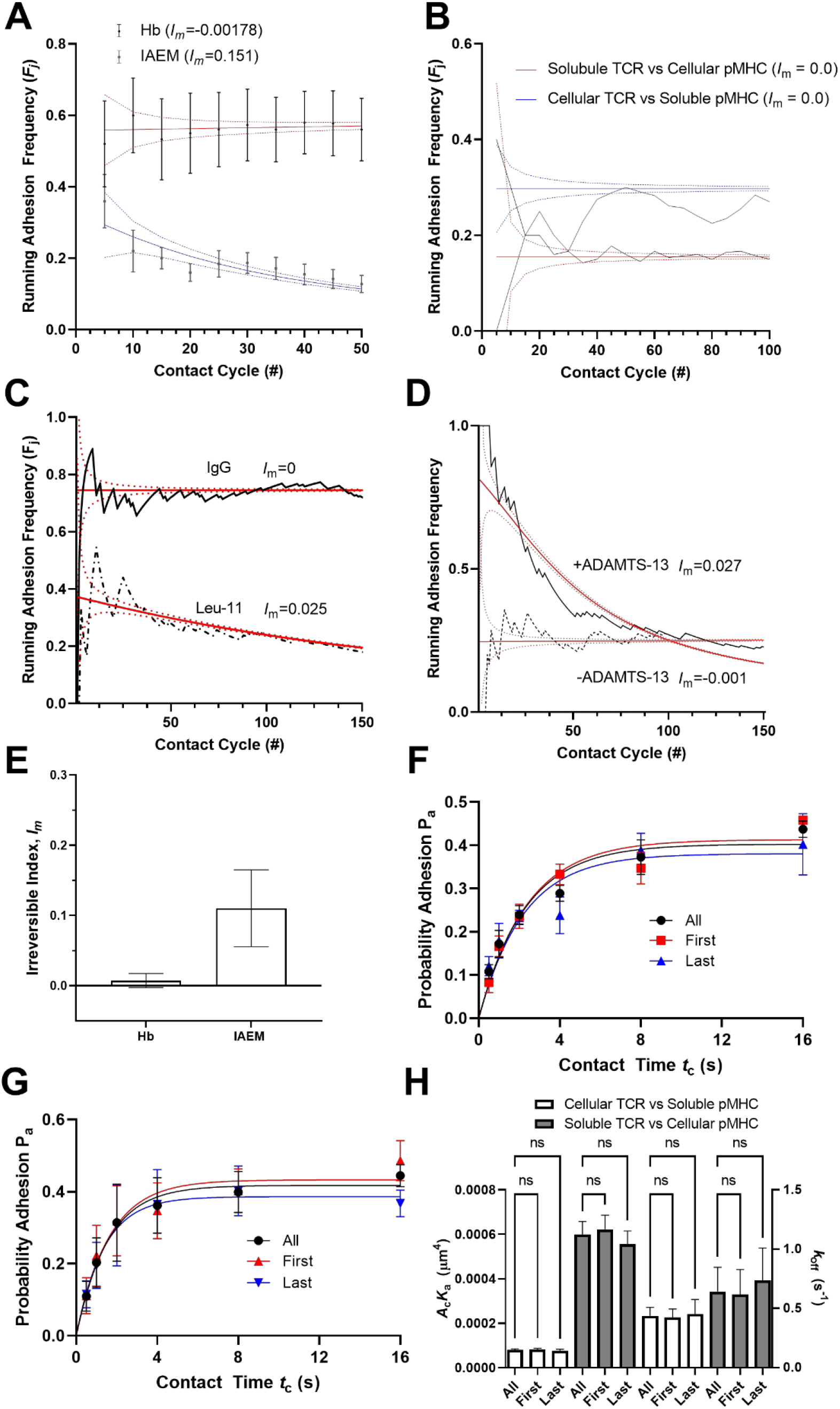
Analysis of memory effect of an intermediate timescale. *A-D*. Representatives of mean ± SEM of running adhesion frequencies vs. contact-retraction cycles data (points) that compare the stability of WT (Hb) and variant (IAEM) peptides presented by I-E^k^ for their interaction with the 3.L2 TCR (A), E8 TCR–TPI:HLA-DR1 interactions in the normal (cellular TCR vs soluble pMHC) and inverted (cellular pMHC vs. soluble TCR) configurations (B), weak GPI anchor FcγRIIIa interacting with its low affinity ligand IgG Fc and with a high affinity anti-FcγRIIIa antibody (C), and GP1bα–VWF A1A2A3 interaction in the presence or absence of solution ADAMTS13 (D). Eq. 6 was fit to the data to evaluate an *I*_m_ value for each curve. The best-fit mean ± SEM curves were plotted as solid and dashed curves along with the data. *E*. Quantifications of the irreversibility indices *I*_m_ of 3.L2 TCR interactions with a panel of peptides complexed with I-E^k^ estimated from 5 sequences of 50 binary adhesion scores for each peptide. *F, G*. Mean ± SEM (n=3 cell-bead pairs per point) of adhesion frequencies *P*_a_ vs. contact time *t*_c_ data (points) fitted by Eq. 1b (curves) for interactions of E8 TCR on Jurkat cells (*m*_r_, 25 *µ*m^-2^) with TPI:HLA-DR1 on BFP beads (*m*_l_, 200 *µ*m^-2^, F), and E8 TCR on BFP beads (*m*_r_, 58 *µ*m^-2^) with TPI:HLA-DR1 on THP-1 cells (*m*_l_, 12 *µ*m^-2^, G). These *P*_a_ vs. *t*_c_ plots are similar to Fig. 2E except that three *P*_a_ values for each *t*_c_ were estimated from three data groups: 1) the first half, 2) the last half, and 3) the entire sequence of ≥50 adhesion scores generated by testing a single cell-BFP bead pair. *H*. The best-fit values of the effective 2D affinity *A*_c_*K*_a_ (*left ordinate*) and off-rate constant *k*_off_ (*right ordinate*) of the E8 TCR–TPI:HLA-DR1 interactions estimated from the three data groups (indicated) in F (open bars) and G (closed bars), with SEM estimated from the scattering of the data by computing the Hessian matrix of the *χ*^2^ function. Each set of parameter values estimated from the three data groups are statistically indistinguishable.

### Analysis of memory effect of a short timescale

We previously demonstrated that binary adhesion score sequences might not obey the i.i.d. assumption, manifesting as both positive (activating) or negative (inhibiting) memory effects depending on the underlying molecular interaction.^28^ In particular, the OT1 TCR interaction with the agonist peptide OVA presented by the mouse MHC class I H2-K^b^α3A2 exhibited positive memory, quantified by a positive increase (Δ*p*) in the positive outcome probability within the present adhesion test given a positive outcome of the immediate past adhesion test. Thus, the adhesion probability at the current contact-retraction test cycle is *p* + Δ*p* or *p* depending on whether the immediate past test cycle resulted in adhesion or no adhesion.

A signature of the positive memory effect is the consecutive presence of the same adhesion score (either 1 or 0) resulting in continuous increase or decrease in the running adhesion frequency *F*_j_ vs. *j* curve, or a same binary score cluster before changing the trend of *F*_j_ or the adhesion score value, as illustrated in Fig. 3A. The measured P14 system cluster size distribution is illustrated in Fig. 3B together with the fit by our previously published model.^28^ The model fit returned the *p* and Δ*p* values for the P14 TCR–gp33:H2-D^b^α3A2 interaction (Fig. 3B). The Δ*p* values of the OT1 TCR interactions with OVA and R4 peptides presented by H2-K^b^α3A2 evaluated by the same fitting methods are shown in Fig. 3C. The positive Δ*p* for OVA is consistent with our previous result, which we now extend to R4 (Fig. 3C). Besides these mouse TCRs on CD8^+^ T cells interacting with class I pMHCs, the interactions between the mouse 3.L2 and human E8 TCRs on CD4^+^ T cells with peptides presented by I-E^k^ and HLA-DR1, which are respective mouse and human class II MHCs, also show significant positive Δ*p* values (Fig. 3D and E). Interestingly, the memory index vanished when we used an inverted configuration to test the E8 TCR–pMHC interaction. In the normal configuration, the E8 TCR was expressed on the Jurkat T cell line and tested by soluble p:HLA-DR1 coated on the BFP glass bead surface. In the inverted configuration, the p:HLA-DR1 was expressed on THP-1 cells and tested by soluble E8 TCR ectodomain coated on the BFP glass beads. This finding suggests that the short-term memory in TCR–pMHC interactions requires cellular TCR expression.

In addition to fitting the model to the data, we also used tone-step transition probability definitions to calculate directly from data (see Fig. 3A) the short-term memory index Δ*p* for the P14 TCR–gp33:H2-D^b^α3A2 interaction^28^ (See Methods). Importantly, the Δ*p* (and *p*) evaluated by the fitting method and the direct calculation agree well in general, as shown in the scattered plots of the values obtained by model fitting vs. the values obtained by direct calculation, which line up along the 45° diagonal line for both Δ*p* (Fig. 3F) and *p* (Fig. 3G), attesting to our methods’ reliability.

Using the non-zero Δ*p* values for the 3.L2 TCR–p:I-E^k^ interactions in Fig. 3D, we can now correct the 2D affinity values estimated using the model of no memory. From Eq. 4, *P*_a_ ≈ *p*/(1 – Δ*p*). From Eq. 1b, *A*_c_*K*_a_ = – ln[1 – *p*(∞)]/*m*_r_*m*_l_ ≈ –ln[1 – (1 – Δ*p*)*P*_a_(∞)]/*m*_r_*m*_l_, indicating that the no memory assumption overestimates the effective 2D affinity. The corrected effective 2D affinity values are plotted along-side with the uncorrected values for comparison (Fig. 2F).

### Analysis of memory effect of an intermediate timescale

In addition to short-term memory effect giving rise to adhesion event clusters but not changing the end running adhesion frequency level, we sometimes observe adhesion frequency sequences in which the *P*_a_ severely underestimates the adhesion probability *p*, as exemplified in Fig. 4A-D. Four cases are shown: 1) 3.L2 TCR interacting with Hb or IAEM presented by I-E^k^ (Fig. 4A), 2) cellular or soluble E8 TCR interacting with soluble or cellular TPI:HLA-DR1, respectively (Fig. 4B), 3) FcγRIIIa interacting with antibodies (Fig. 4C), and 4) GP1bα interacting with VWF A1A2A3 tri-domain in the presence of an ADAMTS-13 solution (Fig. 4D).

In the first case, the IAEM peptide is a mutant form of the WT peptide Hb for the 3.L2 TCR with residue substitutions greatly reducing peptide MHC-II molecule interaction.^36^ The weakened anchor might cause the IAEM peptide to dissociate and be replaced by the null peptide MCC (see Methods) causing ligand function loss. This did not occur for the WT peptide Hb as it was stably bound to MHC (Fig. 4A). A lack of intermediate memory was observed in the second case of E8 TCR interacting with TPI:HLA-DR1 regardless of whether the system was tested in the normal (cellular TCR vs. soluble pMHC) or inverted (soluble TCR vs. cellular pMHC) configuration (Fig. 4B). In the third case, CHO cells expressing FcγRIIIa-GPI were tested by RBCs coated with human IgG or an anti-FcγRIIIa mAb (Leu-11).^10^ The GPI membrane isoform is a fusion protein where the transmembrane and cytoplasmic segments of the WT FcγRIIIa have been replaced by a glycosylphosphatidylinositol (GPI) C-terminus for outer leaflet plasma membrane molecule mounting.^10^ During the retraction phase of the contact-retraction cycles, the FcγRIIIa-GPI molecule might be uprooted from the cell membrane because the GPI anchor is thought to be weaker than the antigen–antibody bond.^34^ In the fourth case, GP1bα on the platelet membrane binds the A1 domain of the plasma protein VWF to initiate the hemostatic and thrombotic cascade.^18^ VWF uses the A3 domain interaction with collagen to immobilize on subendothelial surface of disrupted blood vessel wall. The adhesiveness depends on VWF multimer size, which is regulated by the plasma proteolytic enzyme A Distintegrin And Metalloproteinase with a ThromboSpondin type 1 motif, member 13 (ADAMTS13) that cleaves a A2 domain peptide bond. This cleavage site is cryptic because it is buried inside the folded A2 domain. Mechanical force on the VWF unfolds the A2 domain, enabling ADAMTS13 to access the exposed cryptic site for proteolytic cleavage, breaking the VWF into smaller multimers.^31^ We previously used the AFM (Fig. 1C) to demonstrate this process of pulling-induced unfolding and resulting cleavage by ADAMTS13 of the A2 domain *in vitro*.^31^

Whereas the biological mechanisms differ, a common feature of the aforementioned processes can be revealed by another non-Bernoulli behavior of the binary adhesion score sequences. This feature manifests as a gradual change in the running adhesion frequency as the number of repeated test cycles becomes larger and larger, as illustrated in Fig. 4A-D. Since a full sequence of test cycles usually takes several minutes to complete experimentally, the memory effect accumulation has an intermediate timescale of minutes. We have developed a phenomenological model for this process type that exhibits a cumulative effect over a progressive running adhesion frequency change, capturing the memory effect by an irreversibility index *I*_m_ ^34^ (see Eq. 6 in Model development). Comparisons of the model prediction and experimental data are illustrated in Fig. 4A-D, showing that our model is indeed capable of fitting the data well in all four cases. The irreversibility indices extracted from 3.L2 TCR interacting with the Hb and IAEM peptides presented by I-E^k^ are shown in Fig. 4E. The different *I*_m_ values can be explained by the long-term memory analysis because the dissociation half times for Hb and IAEM are >100 hrs and <20 min, respectively (see below).

As another way to evaluate PPI memory of the intermediate timescale, we compared adhesion probabilities estimated from different segments of the binary adhesion score sequence measured from the same cell over time. We ran repetitive adhesion tests using multiple cells. For each cell the data were segregated into two groups consisting of adhesion scores measured from the first and last half of the contact cycles to estimate two adhesion frequencies. For each group the adhesion frequencies *P*_a_ at the same contact time *t*_c_ were pooled from all cells tested to generate two *P*_a_ vs. *t*_c_ plots to compare with each other and with all the data without segregation. We generated such *P*_a_ vs. *t*_c_ plots for adhesions mediated by interactions of E8 TCR on Jurkat cells with TPI:HLA-DR1 on BFP beads (Fig. 4F), and E8 TCR on BFP beads with TPI:HLA-DR1 on THP-1 cells (Fig. 4G). In both cases, the effective 2D affinity *A*_c_*K*_a_ and off-rate *k*_off_ evaluated from the three groups of data are statistically indistinguishable (Fig. 4H).

### Analysis of memory effect of a long timescale

Depending on the peptide, the timescale of peptide dissociation from MHC molecules varied from tens of minutes to tens of hours. The latter time is much longer than the time spent measuring a typical sequence of repeated adhesion tests, making it difficult to quantify the long-term memory effect using the irreversibility index *I*_m_. To assess the loss of binding function due to slow dissociation of the peptide from the pMHC complex, we developed an adhesion frequency-based assay where the cognate peptide dissociated from the RBC-coated pMHC molecule in a solution without cognate peptide but with a high null peptide concentration to block dissociated cognate peptide rebinding (see Methods). Using five pairs of T cells and RBCs we performed five sequences of 50 repeated 5-s contact duration adhesion tests for each TCR–pMHC pair. Given the fast off-rates of the TCR–pMHC interactions studied here, 5-s contact duration is long enough (i.e., *t*_c_⟶∞) for the adhesion frequency *P*_a_ to reach steady-state^11, 13^ (Fig. 2D). Although ~30 min was required to measure five pairs of cells, we assumed that the mean ± SEM *P*_a_ values were measured at a single elapsed time point starting from the initial time (*t*_e_ = 0). We then generated two sets of steady-state *P*_a_ vs. elapsed time *t*_e_ data, one for OT1 TCR and the other 3.L2 TCR, each interacting with a panel of its cognate pMHC ligands (Fig. 5A-C). As can be seen, the *P*_a_ for different peptides displayed different rates of decay over time. The data were fitted by Eq. 8d to evaluate the peptide dissociation rate constant, *k*_p_, or the half time for peptide dissociation, *t*_½_ = ln2/*k*_p_. The *t*_½_ values for the two TCRs interacting with their corresponding peptides are plotted in Fig. 5D and E showed wide variations depending on the peptide.

**Figure 5.**
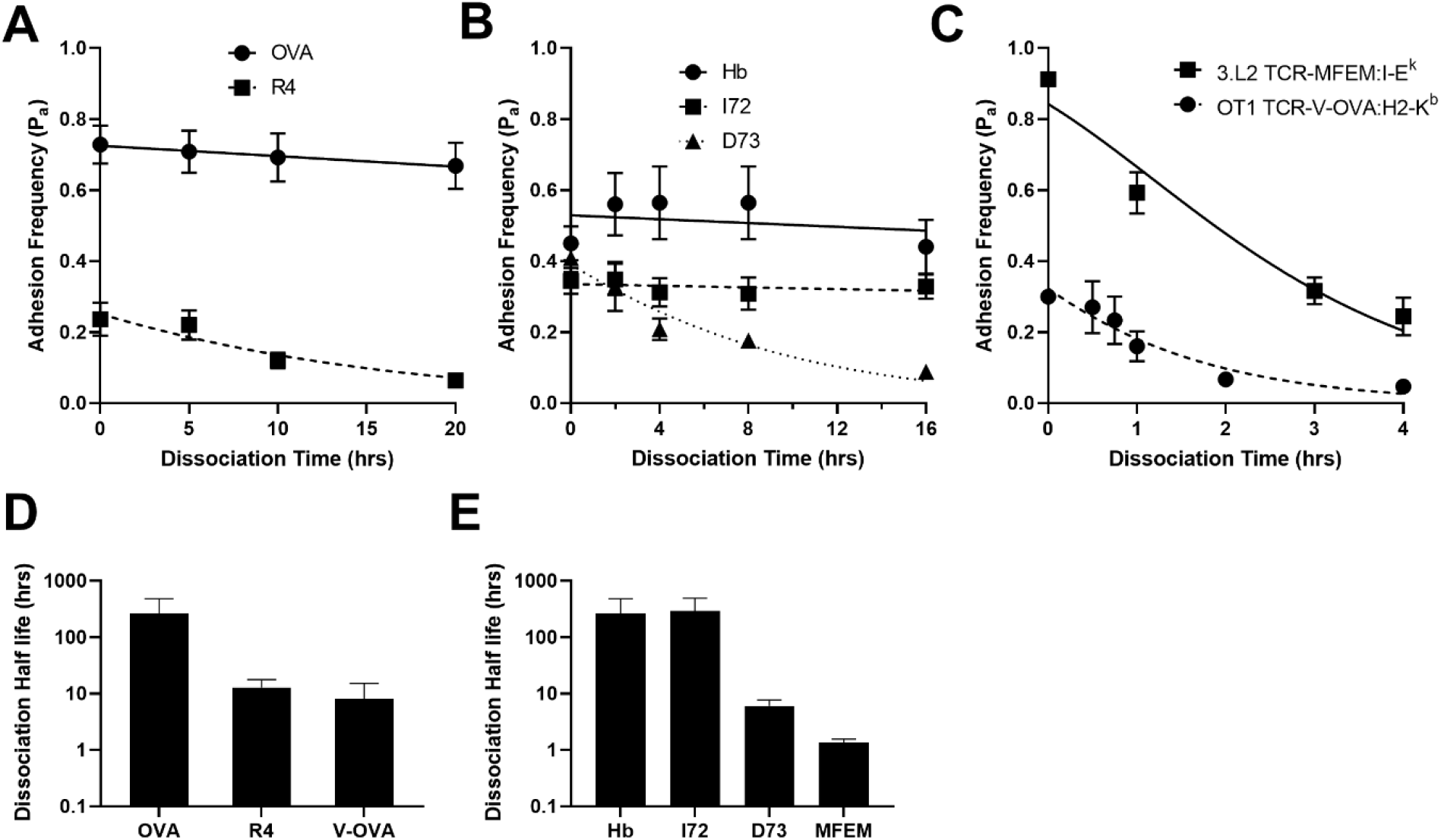
Analysis of memory effect of a long timescale. *A-C*. Mean ± SEM (n ≥ 4) steady-state adhesion frequencies *P*_a_(∞, *t*_e_) of OT1 (A, C) or 3.L2 (B, C) TCR interacting with the indicated pMHC ligands recognized by the specific TCRs, are plotted vs. elapsed time *t*_e_ for peptide dissociation. Data (points) are fitted by Eq. 8d (curves) to evaluate *k*_p_, the rate constant for peptide dissociation. *D, E*. Time required for peptide to dissociate to half of its initial site density, calculated by *t*_½_ = ln2/*k*_p_ using the best-fit *k*_p_ values from A-C, are shown for the OT1 (D) and 3.L2 (E) TCR interacting with the indicated pMHC ligands. Error bar = SEM estimated from the scattering of the data in (A-C) by computing the Hessian matrix of the *χ*^2^ function.

### Interplay between intermediate and long timescales

For a peptide that dissociates at a *t*_½_ of several tens of minutes, the interval between two successive elapsed time points should be no longer than 10 min to obtain enough time points in a dissociation curve for fitting. As such, the ~30 min time required to measure five sequences of 50 repeated adhesion tests each is too long to be approximated by a single time point. We therefore developed a hybrid analysis to bridge the intermediate and long timescales (see Methods). We began the experiment at *t*_e_ = 0 and performed five 50 repeated adhesion tests at 0, 10, 20, 40 and 50 min to generate five *P*_a_ values at these time points using a single pair of cells. This was repeated four times using another four pairs of cells to generate four more mean ± SEM *P*_a_ values at the midpoints of the ~5.8 min period taken to complete each 50 consecutive tests, each right-shifted by 10 min relative to the previous point. We then fitted our *P*_a_ vs. *t*_e_ data with Eq. 8d to evaluate a long timescale memory index, *k*_p_ = 0.04 ± 0.006 s^-1^, and the calculated half-time for peptide dissociation, *t*_½_ = 17.4 ± 0.7 s (Fig. 6A).

**Figure 6.**
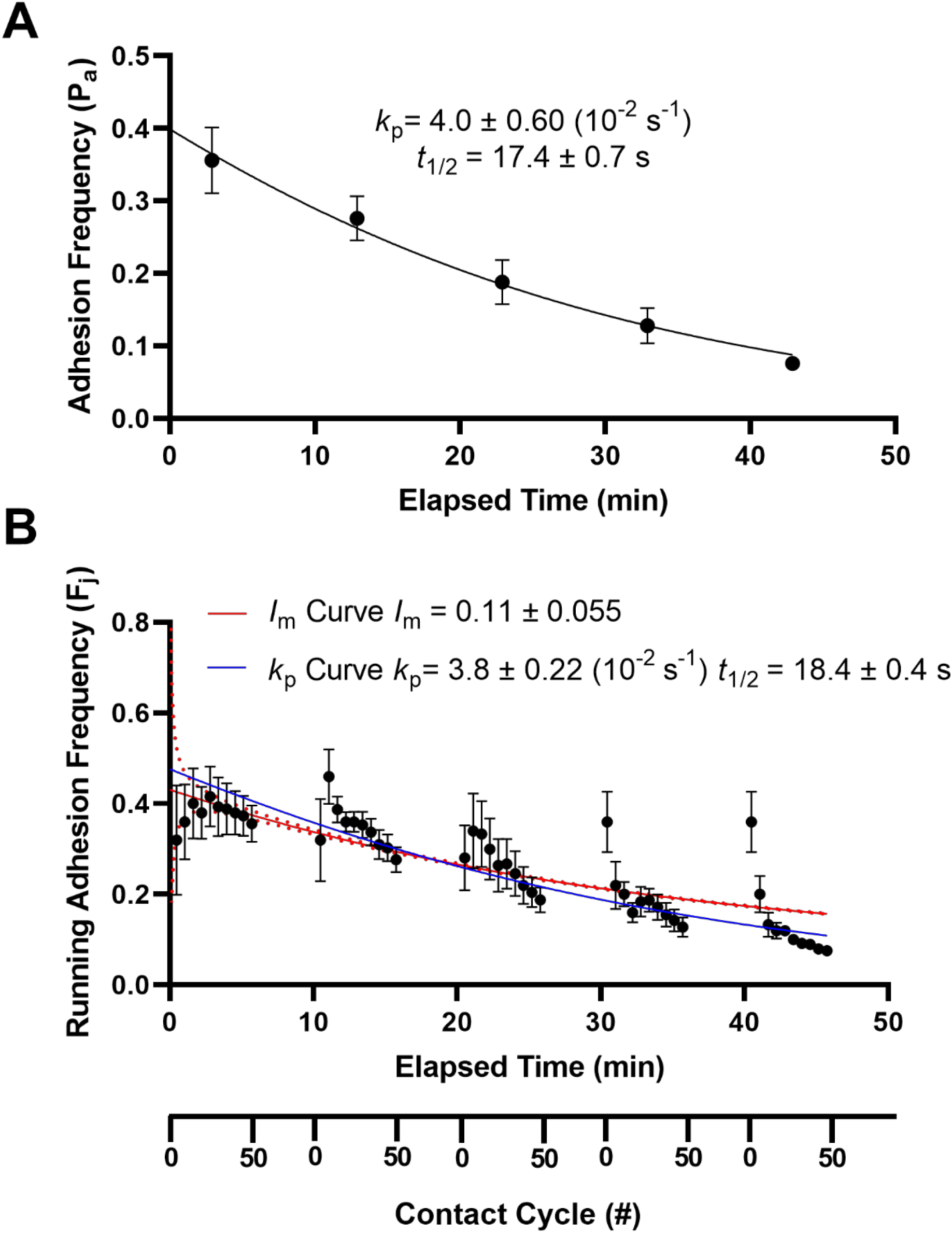
Interplay between intermediate and long timescales. *A*. Mean ± SEM (n = 5 pairs of cells) of adhesion frequency vs. elapsed time *t*_e_ for the 3.L2 TCR–IAEM:I-E^k^ interaction. The 50 repeated adhesion tests started at = 0, 10, 20, 20, and 40 min but took ~5.8 min to complete because each test cycle took ~7 s. We therefore place the lump data point at the middle of the 5.8 min duration of data acquisition. Data (points) were fitted by Eq. 8d to evaluate the peptide dissociation rate constant *k*_p_. *B*. Alternative analysis by expanding the lump data in (A). Mean ± SEM of running adhesion frequency vs. elapsed time *t*_e_ (1^st^ *x*-axis) or contact-retraction cycles (1^st^ *x*-axis). Data were fitted by both Eq. 8 to evaluate a peptide dissociation rate constant *k*_p_ and Eq. 6 to evaluate the irreversibility index *I*_m_. The half times of peptide dissociation calculated by *t*_½_ = ln2/*k*_p_ using the best-fit *k*_p_ values are indicated in A and B, which are statistically indistinguishable.

Alternatively, we plotted the mean ± SEM running adhesion frequency *F*_j_ of the five 50 contact cycle sequences vs. test cycle number *j* (2_nd_ *x*-axis) and elapsed time (1_st_ *x*-axis) in the same graph (Fig. 6B) but the starting point of each sequence was right-shifted by 10 min. These data were fitted to Eq. 8d to evaluate another *k*_p_ (= 0.038 ± 0.0022 s^-1^) and *t*_½_ (= 18.4 ± 0.4 s) statistically indistinguishable from the previous value obtained from the first approach, shown by their comparable peptide dissociation half times (Fig. 6A and B).

The data in Fig. 6B were also fitted by Eq. 6 to evaluate an *I*_m_, the memory index of the intermediate timescale. The best-fit *F*_j_ vs. *j* curve plotted in Fig. 6B was similar to the best-fit *P*_a_ vs. *t*_e_ curve. The *t*_½_ = 24 min evaluated from the dissociation analysis corresponds to 214 contact cycles and an *I*_m_ value of 0.0134.

### Memory analyses of force-dependent bond lifetimes

As mentioned earlier, the original goal of the adhesion frequency assay was to measure *in situ* cross-junctional receptor–ligand interaction kinetics on living cell surfaces.^1, 6, 7^ In the no memory, short-term memory, intermediate-term memory, and long-term memory cases, the 2D kinetic parameters could be calculated from *p* using Eqs. 1, 5, 6, and 8, respectively. In the latter three cases, the memory effects in the short, intermediate, and long timescales were quantified by Δ*p, I*_m_, and *k*_p_, respectively. Note that the effective 2D affinity *A*_c_*K*_a_ estimated from adhesion frequency *P*_a_ had to be corrected in the presence of a nonzero Δ*p* (Fig. 2E). Since affinity is the ratio of on-rate over off-rate, we asked the question of whether the presence of memory effect impacts on-rate, off-rate, or both.

To answer this question, we used the BFP force-clamp assay to measure single receptor– ligand bond lifetimes under a range of constant forces.^11, 13, 30, 35, 37^ Like the adhesion frequency assay, the force-clamp assay also generates a time series from each cell tested repeatedly. Unlike the adhesion frequency assay that generates two types of events (binding and no-binding), the time series resulted from the force-clamp assay includes three types of events: no-binding events, binding events, and lifetime events (Fig. 7A). Like the adhesion frequency assay, the first two types of events can be respectively quantified by binary scores of 0’s and 1’s. Unlike the adhesion frequency assay, the last type of events must be quantified by a continuous positive variable of bond lifetime *t*_b_ (in s). The reciprocal average bond lifetime is equal to off-rate, i.e., <*t*_b_> = 1/*k*_off_ for a first-order irreversible dissociation.^4^ A short-term memory index can be defined similar to the definition of Δ*p* but quantifies memory using a differential bond lifetime Δ<*t*_b_*>*. This can be done by letting the average of bond lifetimes measured from lifetime events following a no-binding event be <*t*_b_*>*, the average of bond lifetimes measured from lifetime events following a binding event be <*t*_b_*> +* Δ<*t*_b_*>*_1_, and the average of bond lifetimes measured from lifetime events following a lifetime event be <*t*_b_*> +* Δ<*t*_b_*>*_2_ (Fig. 7A). To evaluate Δ<*t*_b_*>*_1_ and Δ<*t*_b_*>*_2_ for the P14 TCR– gp33:H2-D^b^α3A2 interaction, all bond lifetime measurements were segregated into three groups: 1) those measured from lifetime events following a no-binding event (group 1), a binding event (group 2), and a lifetime event (group 3). Unsegregated and segregated bond lifetime measurements were binned according to the forces under which they were measured, averaged, and plotted vs. force. It is evident from Fig. 7B and C that the average bond lifetime vs. force curves of all four cases are statistically indistinguishable, indicating the lack of short-term memory in the P14 TCR–gp33:H2-D^b^α3A2 bond lifetimes. Since the adhesion frequency test sequences contain a positive Δ*p* (Fig. 3E), our results suggest that short-term memory only impacts on-rate but not off-rate for P14 TCR–gp33:H2-D^b^α3A2 molecular interactions.

**Figure 7.**
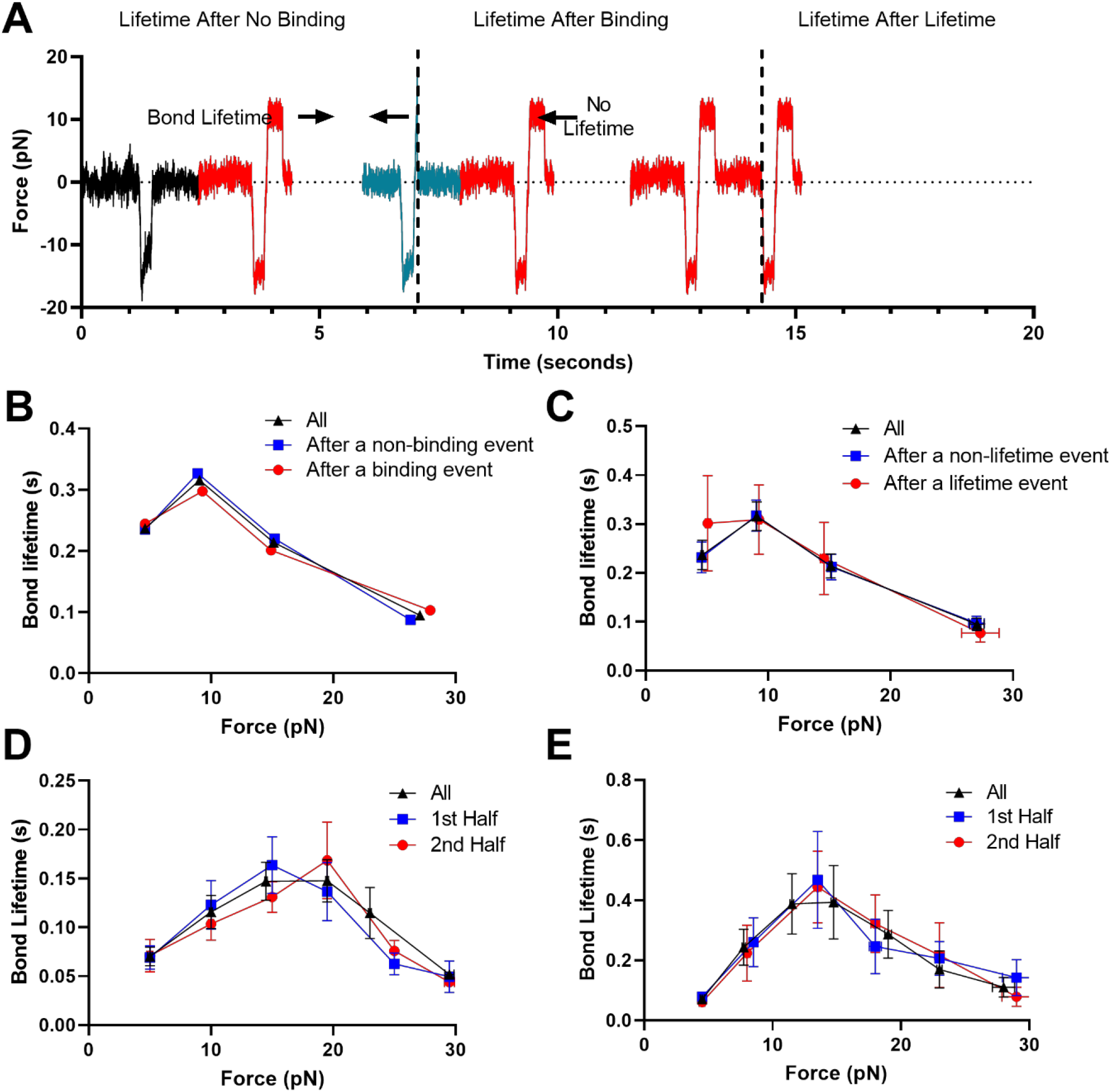
Memory analyses of force-dependent bond lifetimes. *A*. Representative force vs. time signals of repetitive force-clamp cycles over a 15.13-s period. Cycles produced different results are color-coded (black: no binding; blue: binding-rupture; red: lifetime). *B* and *C*. Mean ± SEM (443 ≥ n ≥ 9 measurements per point) of lifetimes of single P14 TCR–gp33:H2-D^b^α3A2 bonds measured after a no-binding event (blue square), after a binding event (red circle), and all events (black triangle) (B), as well as after a non-lifetime event (blue square), after a lifetime event (red circle), and all events (black triangle) (C). *D* and *E*. Mean ± SEM (199 ≥ n ≥ 9 measurements per point) of single bond lifetimes of full E8 TCR-CD3 complex expressed on Jurkat cells interacting with TPI:HLA-DR1 coated on BFP beads (D) or soluble E8 TCRαβ ectodomain coated on BFP beads interacting with membrane TPI:HLA-DR1 expressed on THP-1 cells (E) measured from the first half (blue square), last half (red circle), and all (black triangle) lifetime events in the time series generated by repetitively testing a single cell.

As a first approximation towards estimating the impact of intermediate memory on off-rate, we asked whether repeated contact cycles on the same T cell would result in changing bond lifetime over time or not. To answer this question, TCR–pMHC bond lifetimes were measured from multiple T cells each repeatedly tested 500-1000 contact cycles, resulting in hundreds of bond lifetime measurements. For each cell the data were segregated into two groups, consisting of bond lifetimes measured from the first and last half of the contact cycles. For each group the bond lifetimes were pooled from all cells tested and plotted vs. force, which are compared with each other and with all the data without segregation. Two sets of experiments were performed: 1) purified E8 TCRαβ ectodomain proteins coated on BFP glass beads interacting with TPI:HLA-DR1 expressing on THP-1 cells, and 2) purified TPI:HLA-DR1 ectodomain proteins coated on BFP glass beads interacting with TCR expressing on Jurkat T cells. Interestingly, the average bond lifetime vs. force curves were statistically indistinguishable in both data sets, suggesting the lack impact of intermediate memory on off-rates and isolating the impact on on-rates.

## Discussion

In this paper, we developed, tested, and/or applied three mathematical models that describe the non-Bernoulli behaviors respectively exhibit in random binary score sequences generated by the adhesion frequency assay^1, 6, 7^ and continuous time series generated by the force-clamp assay.^11, 13, 30, 35, 37^ These two assays are designed to mechanically measure the *in situ* reaction kinetics of receptor–ligand interactions in the absence and presence of force, respectively. The assays use a force transducer functionalized with specific ligands to repeatedly contact a living cell expressing, or a surface functionalized with, receptors of interest. By taking the advantage of the ultrasensitivity of the force transducers used in this assay, including MP, BFP, and AFM, which can detect the formation of as low as a single receptor–ligand bond, the experimenter can minimize the living cell measurement disturbances. However, receptor–ligand interactions at the level of a low number of bonds are stochastic. Despite the best effort of the experimenter to ensure similarity between two contact-retraction cycles, the test outcomes may not be the same because they are random. With both assays, whether adhesion would occur in any single test is probabilistic. With the force-clamp assay, whether a binding event would survive ramping force to generate a lifetime event is probabilistic, and lifetime of how long a particular bond would last is also randomly distributed. For these reasons, many tests are required to estimate the probabilities or probability distribution. One approach is to simultaneously measure a large number of cells in parallel, such as those done using a centrifugation assay^38, 39^, a cell collision assay^40, 41^ or a resetting assay ^42^. A more commonly used approach tests one cell at a time but repeats the test many times in series. This is the approach of the adhesion frequency assay and force-clamp assay, which generate a random sequence of binary adhesion scores and a random series of bond lifetimes, respectively.

To extract receptor–ligand binding kinetics information from the random sequence of binary adhesion scores requires two steps: 1) estimate the probability of adhesion and 2) relate the adhesion probability to the kinetic parameters and other experimentally controlled variables. For the second step, we developed a probabilistic equivalent of the well-established deterministic reaction kinetics model by replacing the bond density with the bond probability distributions^1, 6^ (Eq. 1). For the first step, our initial publication assumed repeated contact-retraction test cycle sequences could be modeled as a Bernoulli sequence.^1^ Bernoulli sequences assume i.i.d. and neglect influences of the past on the present, thereby neglecting memory to simplify the analysis. Nearly a decade after the first publication of the adhesion frequency assay, we reported an improved treatment of the random sequence of binary adhesion scores by modeling it as a one-step Markov process without the i.i.d. assumption.^28, 32, 33^ By taking into consideration the influences of the past on the present, which we termed here as memory of a short timescale (seconds), we extracted additional information, the memory index Δ*p* from the binary time sequences.

The present work first extended the application of the Markovian model to three TCRs interacting with their corresponding panels of pMHCs of class I or II, which further confirmed model validity and demonstrated that short-term memory is not an isolated phenomenon but commonly observable in TCR–pMHC interactions. Importantly, we showed that this memory requires the full-length TCR-CD3 complex to reside on the T cell, as memory vanished when soluble TCRαβ ectodomain was tested. Also, the 2D affinity estimated from the adhesion frequency assay should be modified to take into consideration the memory effect without which the *A*_c_*K*_a_ value would be overestimated (for the case of positive Δ*p*).

The main part of this paper was the development, validation, and application of two new models for analysis of the random sequences of binary adhesion scores to capture non-Bernoulli behaviors in intermediate (minutes) and long (hours) timescales. The model of intermediate timescale considers the cumulative memory effect of the entire past event sequence, not just the immediate past of the current event as in the case of the one-step Markovian model for the short timescale memory. Unlike the short timescale memory, the underlying biological mechanisms remain to be elucidated, several mechanisms have been suggested to give rise to memory effects in the intermediate and/or long timescales. These include receptor up- or down-regulation due to cell activation, inhibition or trafficking, receptor extraction from the cell membrane by mechanical force, ligand instability due to peptide dissociation, ligand fragmentation by proteolytic cleavage, etc. By extending the original adhesion frequency assay, we presented examples of some of these cases to demonstrate the universal applicability of our models. Not only do our models help organize the data and provide more accurate data, but they also extract new information that otherwise unobtainable, including the irreversibility index *I*_m_ and the peptide dissociation rate constant *k*_p_ or the half time for peptide dissociation.

We generalized the short-term memory metric from Δ*p* measured by the adhesion frequency assay to Δ<*n>*_1_ and Δ<*n>*_2_ measured by force-clamp assay. Regardless of the metrices and how they are quantified, the concept is the same, i.e., whether and how a past interaction influences the present interaction. The generalization lies in the parameter used to characterize interaction. It is necessary because we model receptor–ligand interaction by reaction kinetics, which requires at least two parameters for its characterization; therefore, two types of memory metrices are required to determine which kinetic parameter, or both, are exhibit memory effects. Our data suggest that for TCR–pMHC interactions, only the on-rate, but not the off-rate, exhibits short-term memory. This finding, together with the finding that only cell surface TCR, but not soluble TCR, exhibits short-term memory, indicate that Δ*p* is underlain by cellular mechanisms rather than the TCR’s molecular properties.

By segregating the time series of random outcomes generated by testing a single cell in a large number of repeated cycles into subgroups depending on when they were measured in the time series (e.g., the first half vs. the last half) and comparing the adhesion frequencies and bond lifetimes measured from the different subgroups, we quantified intermediate memory in a model-independent way. Applying this analysis to the TCR–pMHC interactions allowed us to test whether the kinetic parameters measured by the adhesion frequency and force-clamp assays are convoluted by cellular changes induced by repeated cycles of serially engaging and exerting force on TCRs on the same cell over time. Our data indicate that there are no detectable changes in the force-dependent TCR–pMHC kinetic parameters within the intermediate timescale during which we tested a single cell, despite that T cells are known to be triggered to signal by such repeated test cycles ^29, 30^. Future studies will extend these models and methods to new applications for different biological systems, relate the memory effects to the biological functionalities of specific cells and specific receptors, and elucidate different biological mechanisms that can give rise to memory effects.

## Materials and Methods

### Mice primary T cells

Transgenic P14 TCR, OT1 and 3.L2 TCR mice were housed at the Emory University Department of Animal Resources facility and experiments followed guidelines of the National Institutes of Health and protocols approved by the Institutional Animal Care and Use Committee of Emory University. Naive T cells were purified via magnetic negative selection from spleens of 6–8 week old mice using either CD4^+^ (for 3.L2) or CD8^+^ (for P14 and OT1) T-cell isolation kit (Miltenyi Biotec) according to the manufacturer’s instructions. Cells were washed and stored at 4°C for up to 24 hrs in R10 media, which consists of RPMI 1640 (Cellgro) supplemented with 10% fetal bovine serum (FBS, Cellgro), 2 mM L-glutamine (Cellgro), 0.01M HEPES buffer (Cellgro), 100 μg/ml gentamicin (Cellgro), and 20 μM 2-β-mercaptoethanol (2-BM) (Sigma-Aldrich).

### Human primary cells and cell lines

Human red blood cells (RBCs) were isolated from the whole blood of healthy volunteers according to a protocol approved by the Institutional Review Board of the Georgia Institute of Technology. For adhesion frequency assay, RBCs were purified by Histopaque-1077, washed with ice cold PBS, and resuspended in EAS-45 buffer (2 mM Adenine, 110 mM D-glucose, 55 mM D-Mannitol, 50 mM Sodium Chloride, 20 mM Sodium Phosphate, and 10 mM L-glutamine). Equal aliquots of RBCs were then mixed with various concentrations of EZ-Link Sulfo-NHS-LC-Biotin (Thermo Scientific) at a pH of 7.2 for 30 min at room temperature, yielding different densities of biotin sites on RBC surfaces. Biotinylated RBCs were washed with EAS-45 buffer and stored at 4°C. For BFP experiments, freshly isolated human RBCs were biotinylated with biotin-PEG3500-NHS (Jenkem Technology) and then incubated with nystatin in N2 buffer (265.2 mM KCl, 38.8 mM NaCl, 0.94 mM KH2PO4, 4.74 mM Na2HPO4, and 27 mM sucrose; pH 7.2 at 588 mOsm) for 30 min on ice. Nystatin-treated biotinylated RBCs were washed twice with N2 buffer and stored at 4°C for BFP experiments.

TCR β-chain deficient Jurkat J.RT3 cells were purchased from ATCC (Manassas, VA) and cultured in RPMI 1640 supplemented with 10% FBS, 100 U/mL penicillin, 100 μg/mL streptomycin, 2 mM L-glutamine, and 20 mM HEPES. J.RT3 were transduced by lentivirus to express the E8 TCR. Briefly, E8 TCR α and β chains joined by a P2A element were subcloned into pLenti6.3 vector with a T2A-rat CD2 reporter. Lentivirus encoding E8 TCR were produced by co-transfection of HEK 293T cells with E8TCR-pLenti6.3, pMD2.G (Addgene #12259), and psPAX2 (Addgene #12260) using lipofectamine 3000 (ThermoFisher Scientific). J.RT3 cells were transduced by incubating overnight with supernatant containing lentivirus and FACS sorted using Aria cell sorter (BD Biosciences) based on the surface expression E8 TCR.

THP-1 cells from ATCC were cultured in RPMI 1640 supplemented with 10% FBS, 100 U/mL penicillin, 100 μg/mL streptomycin, 2 mM L-glutamine, and 20 mM HEPES. One day prior to experiment, the cells were treated with 20 – 100 U/ml IFN-γ (R&D Systems) and 1 μM TPI peptide (Genscript) for surface expression of TPI:HLA-DR1 ligand for the E8 TCR.^35^

### Proteins and antibodies

Biotinylated E8 TCRαβ were prepared as previously described.^43^ Briefly, a 17-amino acid tag (TPI, GGGLNDIFEAQKIEWHE) was added to the C-terminus of the E8 TCRα ectodomain. E8 TCRαβ ectodomain protein was produced by *in vitro* folding from inclusion bodies expressed in *E. coli*.^19^ The purified proteins were biotinylated using biotin protein ligase (Avidity); excess biotin and ligase were removed with a Superdex 200 column (GE Healthcare).

The following recombinant pMHC class I and II monomers were from the National Institutes of Health Tetramer Core Facility at Emory University. For P14 TCR, the ligand was the LCMV Armstrong peptide gp_33-41_ (KAVYNFATM) presented by H2-D^b^ (mouse MHC-I) with a C-terminal biotin tag.^44^ For the OT1 TCR, the ligands consisted of the wild-type (WT) peptide OVA^323-339^ (SIINFEKL) or an altered peptide R4 (SIIRFEKL) or V-OVA (RGYNYEKY), or a control peptide VSV (RGYVYQGL) presented by H2-K^b^ (mouse MHC-I) with a C-terminal biotin tag ^13^. Both mouse MHCs were mutated (MT) by replacing their α3 domain by that of the human HLA-A2 to prevent CD8 binding (H2-D^b^α3A2 and H2-K^b^α3A2). For the E8 TCR, the ligand was the melanoma antigenic peptide TPI derived from the glycolytic enzyme triosephosphate isomerase (GELIGTLNAAKVPAD) presented by HLA-DR1 (human MHC-II) with a C-terminal biotin tag ^35^.

For the 3.L2 TCR, I-E^k^ (mouse MHC-II), a generous gift from Peter Jensen (Emory University), was initially bound by a CLIP peptide but replaced prior to experiment by either the WT peptide murine hemoglobulin epitope Hb_64-76_ (GKKVITAFNEGLK), or an altered peptide I72 (GKKVITAFIEGLK) or D73 (GKKVITAFNDGLK), or IAEM (GKKVITAAIEGLM) or MFEM (GKKVMTAFNEGLM), or a control peptide MCC from the moth cytochrome c (ANERADLIAYLKQATK).^36^ The latter two peptides were produced by replacing two or three MHC anchor residues 68I, 71F, 73E and 78K of the Hb sequence at positions 1, 4, 6, and 9 by M, A, P, or M.^36^ The null peptide MAEM (GKKVMTAANEGLM) was used as negative control.^36^

The extracellular fragment of GPIbα (glycocalicin)^45^ was purified from platelets as described.^46^ Recombinant A1A2A3 with a 6-histidine tag at the C-terminus^47^ and ADAMTS-13^48^ were generous gifts of Miguel Cruz and Jinf-fei Dong, respectively (Baylor College of Medicine) and produced as previously described. Anti-His mAb was from Sigma (St. Louis, MO). Anti-A1 mAb CR1 was a generous gift from M. C. Berndt (University College Cork, Cork, Ireland).

### Coating proteins on surfaces

Two methods were used to coat pMHC on RBCs for the micropipette experiment – biotin– streptavidin (SA) coupling (for p:H2-K^b^) and chromium chloride (CrCl_3_) coupling (for p:I-E^k^) the detailed procedures of both of which have been described previously.^1, 13^ Briefly, RBCs were biotinylated, conjugated with streptavidin, and incubated with C-terminally biotinylated p:H2-K^b^.^13^ Alternatively, RBC’s in 4% hematocrit were suspended in a 0.001% solution of CrCl_3_ in 0.02M acetate buffer, pH 5.5. When 10 μg/ml p:I-E^k^ in phosphate free media was added, spontaneous coupling occurred. After 5 min the reaction was quenched with PBS, 5 mM EDTA with 1% BSA. RBC’s were used immediately after protein coating.

To coat TCR or pMHC on glass beads for BFP experiments, silanized beads were covalently linked to SA-maleimide (Sigma-Aldrich) then conjugated with subsaturating C-terminally biotinylated TCR or p:MHC.^11, 24, 37^

To functionalize the AFM system, cantilever tips were incubated with 10 µl per tip of GPIbα (10 μg/ml), CR1, (15 μg/ml) or BSA (1%) at 4 °C overnight, rinsed, and soaked in PBS containing 1% BSA to block nonspecific binding. Polystyrene dishes were thoroughly cleaned with absolute ethanol and dried with argon gas before protein adsorption. Surfaces were incubated with 10 µl per spot of A1A2A3 or anti-His mAb (15 μg/ml) at 4 °C overnight and washed two times with PBS. The anti-His mAb coated surfaces were further incubated with 10 µl per spot of A1A2A3 (5 μg/ml) at room temperature for 1 h. Dishes were then filled with PBS containing 1% BSA in the absence or presence of ADAMTS-13 (1.25-10 µg/ml) without or with 5 mM EDTA.

### Site density measurement

Site densities of the TCR and pMHC were measured by flow cytometry^1, 11, 13^ using fluorescent antibodies. Antibodies were used at 10 μg/ml concentration in 100 μl of FACS buffer (PBS without calcium and magnesium, 5 mM EDTA, 1% BSA, 25 mM HEPES, 0.02% sodium azide) at 4 °C for 30 min; fluorescent intensities were measured by the BD LSR II flow cytometer (BD Biosciences); and site densities were calculated by comparing to the BD QuantiBRITE PE standard beads (BD Biosciences).

### Adhesion frequency assay using micropipette (MP), biomembrane force probe (BFP), or atomic force microscopy (AFM)

The detailed procedures of this assay using MP have been described previously^1, 6, 7^ and it was used to test adhesions mediated by OT1 TCR–p:H2-K^b^ and 3.L2 TCR–p:I-E^k^ interactions. Briefly, two micropipettes holding a T-cell on one side and a pMHC-coated RBC on the other (Fig. 1A) were controlled to make repeated contacts with a constant area (*A*_c_). From the presence or absence of RBC shape elongation upon retraction after a contact time (*t*_c_), the adhesion scores (1 for adhesion and 0 for no adhesion) in no less than 50 repeated contact-retraction cycles were enumerated to generate a binary sequence.

The adhesion frequency assay was also performed using BFP or AFM. The only difference was that the RBC was replaced by a pMHC-coated force probe bead (in the case of BFP) or a microcantilever (in the case of AFM). Instead of observing the RBC shape elongation microscopically, the displacement of the bead or the deflection of the cantilever gave rise to a tensile force signal that signified adhesion.

Our BFP system (Fig. 1B) has been described^11, 13, 30, 35, 37^ and it was used to test adhesions mediated by P14 TCR–p:H2-D^b^ interaction.

The AFM (Fig. 1C) was built in our laboratory as previously described.^4, 31^ It was functionalized in two ways: 1) A1A2A3 was adsorbed on a polystyrene surface and tested by an AFM tip coated with glycocalicin and 2) A1A2A3 was captured by an anti-His mAb preadsorbed on a polystyrene dish and tested by an AFM tip coated with the monoclonal antibody (mAb) CR1. To assess the memory effect in the intermediate timescale due to proteolytic cleavage, the experiments were performed with or without ADAMTS13 in the media in the presence or absence of EDTA.

### Peptide dissociation assay

To assess the memory effect of the long timescale due to ligand instability resulted from peptide dissociation from the pMHC, we first incubated the CLIP pMHC-coated RBCs in 100 μM of the peptide to be tested for a prolonged period of time to allow CLIP peptide on all pMHC molecules coated on the RBCs to be replaced by that peptide. At time zero, the RBCs were spun down and the pellet was resuspended in an infinitely dilute solution to allow the peptide to dissociate from pMHC. A high concentration (100 μM) of null peptide (VSV or MCC for the RBCs bearing p:H2-Kb or p:I-E^k^, respectively) was added to the solution to further block rebinding of the dissociated peptide. The clock for elapsed time was then turned from this point on. At each elapsed time point, an aliquot of RBC solution was added to a new cell chamber on top of the microscope stage and the micropipette adhesion frequency assay was performed to measure 5 sequences of 50 adhesion scores each using five T cell–RBC pairs to generate sufficient data for mean ± SEM. For a contact of 5-s duration, it took ~5.8 min to perform 50 repeated contact-retraction tests using a single cell pair and ~30 min to complete the assay with five pairs of cells. The 5-s contact time was chosen because it is long enough for the adhesion frequency *P*_a_ and adhesion probability *p* to reach steady-state (cf. Fig. 2D), which simplifies our analysis because we could take *t*_c_ as infinity in Eq. 8, making it dependent on *t*_e_ only.

For slow dissociating peptides that give rise to memory indices of long timescales (many hours), it seems reasonable to approximate a 30-min period by a single elapsed time point. And the experiment was repeated in different elapsed time points to generate a *P*_a_(∞,*t*_e_) vs. *t*_e_ curve for evaluation of *k*_p_ by fitting Eq. 8d to the data. However, for memory effects of timescale of tens of minutes, which lies in between the intermediate and long timescales by our definitions, a 30 min period seemed too long to be considered as a single elapsed time point. To bridge the two timescales characterized by *I*_m_ and *k*_p_, we designed the following experiment.

Instead of generating five sequences of 50 adhesion scores each using five T cell–RBC pairs upon adding to the cell chamber RBCs that had been in infinitely diluted media for peptide dissociation for a given elapsed time, we used a single RBC to test a single T cell for five sequences of 50 consecutive contact-retraction cycles upon adding to the cell chamber RBCs newly resuspended in infinitely diluted media from the cell pellet, i.e., RBCs at elapsed time zero. The experimenter started the first, second, third, fourth, and fifth 50 consecutive cycles at *t*_e_ = 0, 10, 20, 30, and 40 min. This experiment was repeated five times using five pairs of cells. These data were then analyzed in two ways.

First, each sequence was broken down in to 5 subsequences of 50 binary scores each for calculation of an adhesion frequency *P*_a_ for that subsequence. The *P*_a_ values of the first subsequences of the five cell pairs were used to calculate the mean ± SEM *P*_a_ for the first elapsed time point approximated by the midpoint of the 5.8 min duration taken to complete 50 repeated tests (2.9 min). The mean ± SEM *P*_a_ values of the second, third, fourth, and fifth subsequences of the five cell pairs were also generated the same way for the second (12.9 min), third (22.9 min), fourth (32.9 min), and fifth (42.9 min) elapsed time points. These *P*_a_ vs. *t*_e_ data were then fitted by Eq. 8d to evaluate a *k*_p_, the memory index of the long timescale (Fig. 6A).

Second, the mean ± SEM running adhesion frequency data of the five 50 adhesion score sequences were plotted vs. the elapsed time (1^st^ *x*-axis) and the contact cycle number (2^nd^ x-axis). The contact cycle number is related to the elapsed time by a conversion factor of 1 cycle = 7 seconds. Since 50 contact cycles took only ~5.8 min, the starting time for next sequence has to be right-shifted by ~4.2 min (Fig. 6B). These data were fitted to Eq. 8d to evaluate another *k*_p_ to compare with the previous value evaluated form the first approach. The data in Fig. 6B were also fitted to Eq. 5 to evaluate an *I*_m_, the memory index of the intermediate timescale.

We first calculated the adhesion frequency *P*_a_ (= average adhesion scores) from the measured binary adhesion score sequences. We also calculated the probability of adhesion *p* and the short timescale memory index Δ*p* using both the direct method (Eq. 2) and fitting to the adhesion cluster size distribution (Eq. 3). Fitting Eq. 1a to the *P*_a_ vs. *t*_c_ data allowed us to evaluate the effective 2D affinity *A*_c_*K*_a_ and off-rate *k*_off_. Using Eq. 4 to account for the difference between *P*_a_ and *p* allowed us to correct the *A*_c_*K*_a_ value in the presence of Δ*p* (cf. Fig. 2E). Fitting Eq. 6 to the running adhesion frequency data returned *I*_m_, the memory index for intermediate timescale. Finally, fitting Eq. 8 to adhesion frequency measured at different elapsed time points returned *k*_p_, the memory index for long timescale.

### Force-clamp assay for force-dependent bond lifetimes using biomembrane force probe (BFP)

This assay has been described previously^11, 13, 30, 35, 37^ and it was used to measure force-dependent lifetimes of P14 TCR–p:H2-D^b^ and E8 TCR–TPI:HLA-DR1 bonds. During the experiment, a protein-coated bead (tracking bead) was attached to the apex of a micropipette-aspirated RBC that serves as a spring with pre-adjusted spring constant of 0.1-0.3 pN/nm. For force clamp assay, the tracking bead was repeatedly contacted by a piezo-driven target bead coated with (or target cell expressing) the corresponding binding partner(s). The displacement of tracking bead was monitored with a high-speed camera at >1000 fps and was translated into force by applying the pre-defined spring constant. During separation, bond formation between tracking bead and target bead/cell pulled the tracking bead away from its baseline, manifesting positive force loading on the molecular bond. Bond lifetime was defined as the duration from the start of clamp at the preset force level to bond rupture. Several hundred bond lifetime events were collected and pooled for various clamp forces using multiple bead-bead or bead-cell pairs.

### Computational Methods for Model Fitting

Model fitting primarily used GraphPad Prism version 9.4.1 compiled for Windows 10 for least squares regression except for Δ*p, p*, and *I*_*m*_ which utilized Python 3.9 with the common analysis packages NumPy and SciPy for Limited-memory Broyden-Fletcher-Goldfarb-Shanno bound (L-BFGS-B) fitting with the appropriate fitting constraints.

## Declarations

None of the authors have any conflict of interest to declare.

## Author contributions

CZ, AG and BDE designed and supervised experiments; CZ developed mathematical models; AMR, YZ, H-KC and FJ performed experiments; AMR, SME and CZ analyzed data and prepared the paper.

## Acknowledgements

We thank the late Professor Robert M. Nerem for his inspiration and teaching over the years. We also thank Jenny Ning Jiang, Kaitao Li and Laurel Ann Lawrence for technical assistance, Peter Jensen for providing purified p:I-E^k^ proteins, and Roy Mariuzza for providing the E8 TCR and TPI:HLA-DR1 proteins. We acknowledge Scott E. Chesla, Tao Wu, and Jiangguo Lin for providing their published data in Refs.^10, 31^ for reanalysis and model fitting. We also acknowledge the National Institutes of Health Tetramer Core Facility at Emory University for providing the pMHC molecules. This work was supported by grants from the National Institutes of Health (T32GM008169 and U01CA214354-S1 to AMR, U01CA250040 to CZ, R01AI124680 to AG and CZ) and by a Postdoctoral Fellowship from the National Research Foundation of South Korea (2021R1A6A3A03038382 to H.-K.C.).

## Graphic Abstract

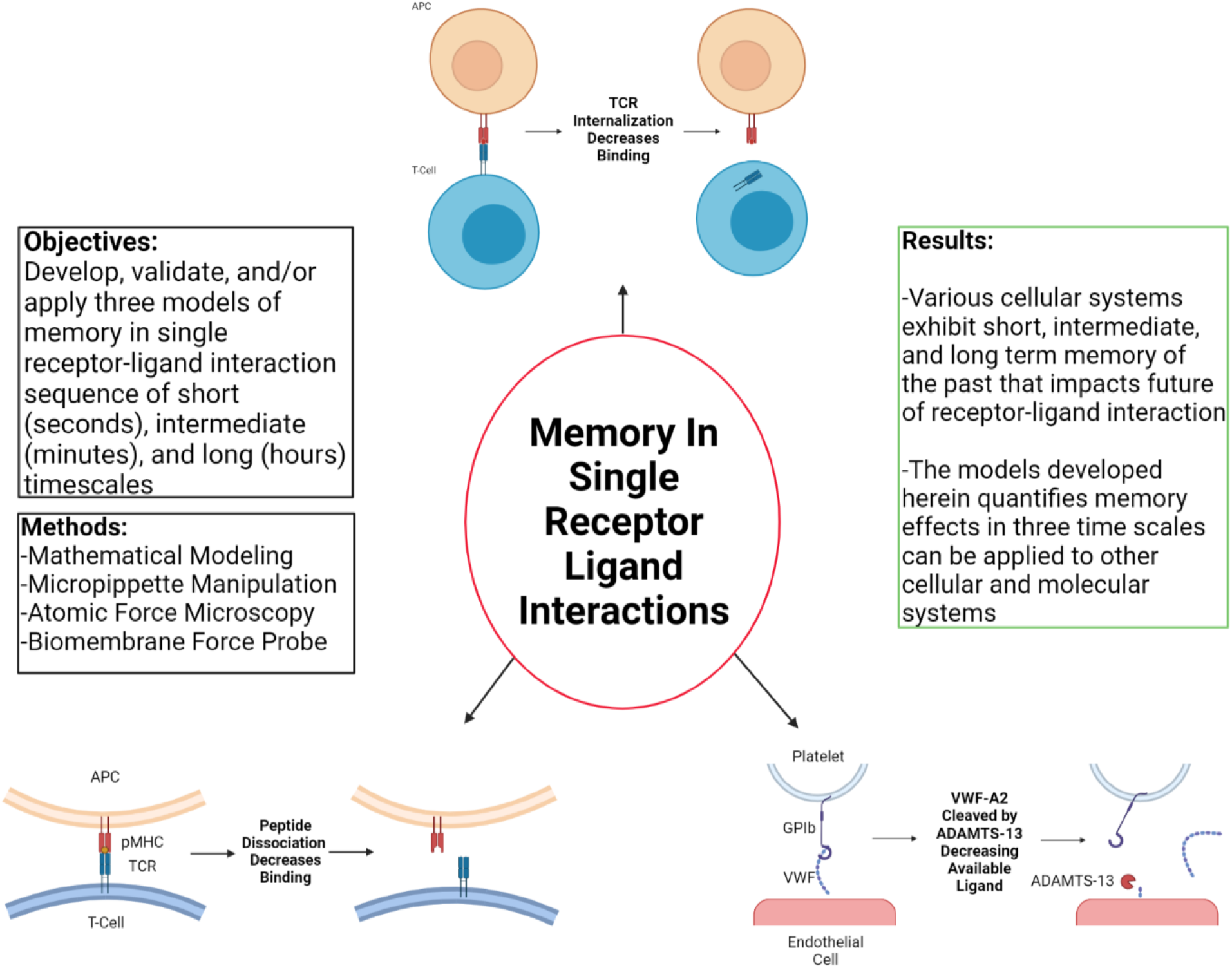

